# 16S rRNA Gene Sequencing as a Clinical Diagnostic Aid for Gastrointestinal-related Conditions

**DOI:** 10.1101/084657

**Authors:** Daniel E. Almonacid, Laurens Kraal, Francisco J. Ossandon, Yelena V. Budovskaya, Juan Pablo Cardenas, Jessica Richman, Zachary S. Apte

**Affiliations:** uBiome, Inc., San Francisco, CA 94105, United States of America; Department of Biochemistry and Biophysics, University of California, San Francisco, San Francisco CA 94158, United States of America

**Keywords:** Next-generation sequencing (NGS), 16S rRNA gene sequencing, microbiome, pathogens, commensal bacteria, clinical microbiology

## Abstract

Accurate detection of the microorganisms underlying gut dysbiosis in the patient is critical to initiate the appropriate treatment. However, most clinical microbiology techniques used to detect gut bacteria were developed over a century ago and rely on culture-based approaches that are often laborious, unreliable, and subjective. Further, culturing does not scale well for multiple targets and detects only a minority of the microorganisms in the human gastrointestinal tract. Here we present a clinical test for gut microorganisms based on targeted sequencing of the prokaryotic 16S rRNA gene. We tested 46 clinical prokaryotic targets in the human gut, 28 of which can be identified by a bioinformatics pipeline that includes sequence analysis and taxonomic annotation. Using microbiome samples from a cohort of 897 healthy individuals, we established a reference range defining clinically relevant relative levels for each of the 28 targets. Our assay accurately quantified all 28 targets and correctly reflected 38/38 verification samples of real and synthetic stool material containing known pathogens. Thus, we have established a new test to interrogate microbiome composition and diversity, which will improve patient diagnosis, treatment and monitoring. More broadly, our test will facilitate epidemiological studies of the microbiome as it relates to overall human health and disease.

## Introduction

Although most microorganisms living within the human host are thought to be harmless or even beneficial [1], some of the deadliest diseases and epidemics in human history have been caused by bacteria [2]. Microbial infections remain a major cause of death and disease worldwide to this day [3]. In addition, the gut microbiome is now recognized as playing a major role in health maintenance and there are clear associations between a microbiome imbalance (dysbiosis) and various diseases and medical conditions [4]. Bacteria in the gastrointestinal tract have been long known to underlie numerous illnesses such as diarrhea and food poisoning. More recently, specific bacteria and associated inflammation have been shown to promote broader gastrointestinal diseases including irritable bowel syndrome [5] and cancer [6].

Rapid and accurate identification of causative microorganisms is critical to provide the appropriate treatment for patients suffering from these gastrointestinal conditions. However, clinical microbiology still largely depends on traditional culturing methods to identify etiological agents. Culture techniques have remained essentially the same for the last 50 years [7]. The specialized work of culturing specific microorganisms is laborious, time-consuming and requires interpretation by extensively trained personnel. Moreover, many organisms are not cultivable and causative agents often fail to grow on culture even when present [8,9]. Alternative techniques to accurately report the composition of the gut microbiome in a timely manner are clearly necessary to improve clinical practice and patient outcome [10].

Next-generation sequencing (NGS) technology [11] can be used to detect prokaryotic 16S rRNA gene sequences within clinical samples, and has the potential to replace culture–based strategies for determining the composition of microbiomes [12,13]. The implementation and consolidation of clinical microbiological tests with NGS would enable rapid, accurate, and reliable detection of all bacteria and archaea present within a sample, including their relative levels, using the 16S rRNA gene sequence as a molecular marker and identifier [4,14,15]. Furthermore, multiplexing allows multiple samples to be processed at the same time, reducing labor and cost, while taxonomic classification can be automated, reducing the need for manual clinical interpretation. Indeed, the identification of prokaryotes by 16S rRNA gene sequencing is in widespread use with many successful examples, including rapid pathogen sequencing [15], microbiome analyses [16-18], and the detection of polymicrobial infections [19]. Compared to traditional clinical microbiology, 16S rRNA sequencing is not only a superior process, but also has a greatly improved sampling procedure (Fig 1). Sequencing requires significantly less material, which eases the patient burden of sample collection, particularly from sources such as fecal matter. Further, the microbial population within the sample is immediately processed by microbial lysing and DNA stabilization, precluding artifacts that might arise after collection. Although ease of sampling is obviously subjacent to accurate detection, it likely facilitates regular sampling for personal health monitoring and public health surveillance.

In contrast to culturing, 16S rRNA gene sequencing can simultaneously detect and quantify numerous bacteria and archaea in a sample. This eliminates the need for physicians to select specific clinical targets for culturing and also creates an opportunity to observe the overall state of the microbiome. This is of particular clinical interest since, concurrent with the advent of NGS, the microbiome has emerged as a major contributor to human health and disease [4]. The gut microbiome especially plays an important role in achieving optimal human health, with demonstrated impacts on aging, metabolic syndrome and immunity. Although medical diagnosis has traditionally focused on pathogens, these overarching and interrelated conditions appear to be greatly influenced by the composition of the commensal gut microbiome. Regularly evaluating the microbiome to monitor overall health is therefore gaining traction in contemporary medicine and needs to be part of modern diagnostics.

In this study, we present the development and validation of a novel NGS-based clinical gut microbiome detection assay. The assay utilizes 16S rRNA gene sequencing to identify 28 clinically relevant microbial targets, 14 species and 14 genera, that comprise pathogens and commensals from the human gastrointestinal tract.

**Fig 1.**
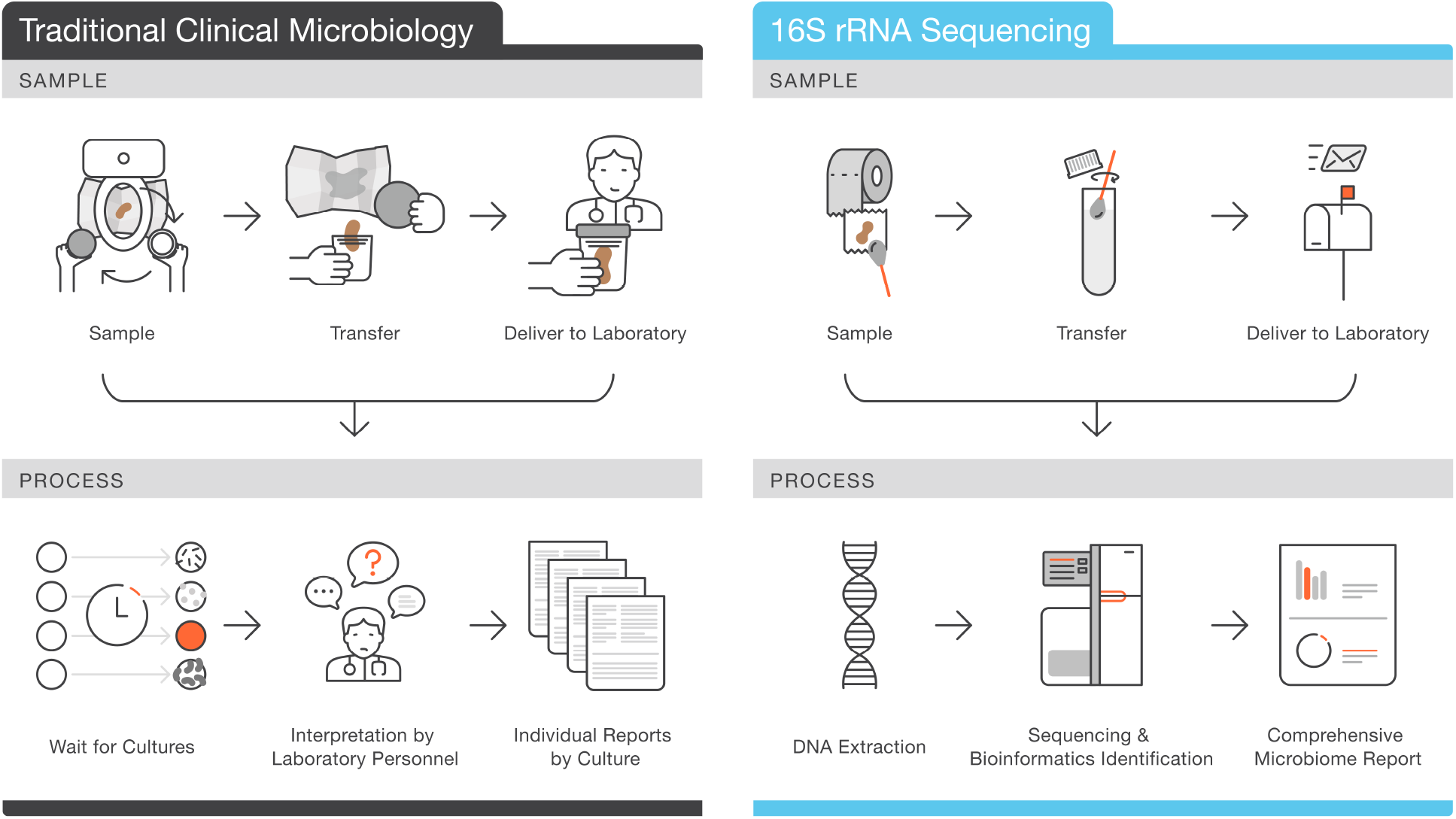
Sample collection and processing of clinical stool samples for traditional clinical microbiology versus 16S rRNA gene sequencing.

A traditional microbiology test performed on a stool sample typically requires collecting a stool sample and immediately delivering it to the laboratory or clinical practitioner. Specific organisms are cultured from the sample based on the physician's requests, followed by laborious and time–consuming processing that requires interpretation by extensively trained laboratory personnel. In contrast, 16S rRNA gene sequencing requires only a fraction of the biological material needed for culture-based techniques (just a swab from toilet paper) and microorganisms are lysed and DNA is stabilized in sample buffer. Thus, sample collection and delivery are greatly simplified. Sequencing and interpretation can be automated to reduce human labor and error.

## Material and Methods

### Sample Collection and 16S rRNA Gene Sequencing

Microbiome samples were collected using commercially available uBiome microbiome sampling kits, which have been designed to follow the specifications laid out by the NIH Human Microbiome Project [20]. Each kit contains lysis and stabilization buffer which effectively kills and lyses all bacteria and archaea and preserves the sample DNA for transport at ambient temperatures. Microbial DNA was extracted in a class 1000 clean room by a column-based approach using a liquid-handling robot. Amplification of the 16S rRNA genes was performed using universal V4 primers (515F: GTGCCAGCMGCCGCGGTAA and 806R: GGACTACHVGGGTWTCTAAT). Library consolidation, column cleanup, size selection and qPCR were performed using standard protocols. Sequencing was performed in a pair-end modality on the Illumina NextSeq 500 platform rendering 2 x 150 bp pair-end sequences. Samples were barcoded with a unique combination of forward and reverse indexes allowing for simultaneous processing of multiple samples.

### Taxonomic Annotation and Reference Database Generation

After sequencing, demultiplexing of samples was performed using Illumina's BCL2FASTQ algorithm. Reads were filtered using an average Q-score > 30. Forward and reverse reads were appended together and clustered using the Swarm algorithm [21] using a distance of 1 nucleotide. The most abundant sequence per cluster was considered the real biological sequence and was assigned the count of all reads in the cluster. The remainder of the reads in a cluster were considered to contain errors as a product of sequencing. The representative reads from all clusters were subjected to chimera removal using the VSEARCH algorithm [22]. Clustered reads passing all above filters were aligned using 100% identity over 100% of the length against a hand-curated database of target 16S rRNA gene sequences derived from version 123 of the SILVA database [23]. The hand-curated databases for each taxa were created by selectively removing sequences with amplicons that were ambiguously annotated to more than one taxonomic group, while still maximizing the sensitivity, specificity, precision and negative predictive value of identification for the remaining amplicons in each taxa (S1 Doc). The relative abundance of each taxa was determined by dividing the count linked to that taxa by the total number of reads passing all filters.

### Participants

Samples from 1,000 self-reported healthy individuals were selected from the ongoing uBiome citizen science microbiome research study (manuscript in preparation). 103 samples did not pass our 10,000 read quality control threshold, resulting in a healthy cohort of 897 samples (62% male and 38% female). Participants were explicitly asked about 42 different medical conditions such as cancer, infections, obesity, chronic health issues and mental health disorders. Selected participants with an average age of 39.7 years (SD = 15.5) responded to an extensive survey and self-reported to be currently and overall in good health. None of the individuals selected for the healthy cohort had ever been diagnosed with high blood sugar, diabetes, gut-related symptoms or any other medical condition. This study was performed under a Human Subjects Protocol provided by an IRB. Informed consent was obtained from all participants. Analysis of participant data was performed in aggregate and anonymously.

### Experimental verification

Double-stranded DNA segments were designed to be representative for the V4 region of the 16S rRNA gene of each target species or genus and synthesized by IDT and Thermo Fisher. Two sets of 14 targets each were combined at 1:10, 1:50, 1:100 and 1:1000 ratios and vice versa, allowing for the detection of different levels of targets in a high background of DNA (the undiluted set). The resulting ratios are 1:14014, 1:1414, 1:714, 1:154 and ~1:14 (1000:14014 for the undiluted set) for each individual target (S1 Fig). The amount of DNA for each target was 1.74 pg (6.078 attomoles) before dilution. Sample combinations were processed in uBiome microbiome sampling kits using the clinical pipeline described above.

Verification samples were obtained from Luminex's xTAG Gastrointestinal Pathogen Panel (xTAG GPP). Verification samples contained real or synthetic stool samples with live or recombinant material, with some specimens being positive for more than one clinical target. 35 positive control samples were used, each certified to be positive for at least one control taxon from our target list, with the exception of those samples containing either *Peptoclostridium difficile* or *Salmonella enterica* which are positive for 2 taxa simultaneously (the species to which they belong and their corresponding genus). The control samples were considered negative for the remainder of the taxa on our test panel. Two out of 35 control samples did not pass our sequencing quality thresholds. Five samples positive for *Yersinia*, a genus that is not present in the final target list, was included as a negative control. Verification samples were processed in uBiome microbiome sampling kits using the clinical pipeline described above.

## Results and Discussion

### Clinically relevant target identification

To derive a preliminary target list of bacteria and archaea to include in our clinical test, we first identified clinically relevant microorganisms present in the human microbiome. We performed an extensive review of the literature and clinical landscape, and obtained evidence supporting the importance of hundreds of microorganisms known to inhabit the human gut. We included these in our initial list, along with organisms that are commonly interrogated in clinical tests. This initial list was further evaluated for positive and negative associations with several indications, including flatulence, bloating, diarrhea, gastroenteritis, indigestion, abdominal pain, constipation, infection, dysbiosis, inflammatory bowel syndrome, ulcerative colitis and Crohn's disease–related conditions. Ultimately, we compiled a preliminary target list containing 15 genera and 31 species of microorganisms associated with human health status (S1 Table), including pathogenic, commensal and probiotic bacteria and archaea.

The bioinformatics annotation pipeline developed for this clinical test was specifically designed to have high prediction performance. To this end, we implemented a taxonomy annotation based on sequence searches of 100% identity over the entire length of the 16S rRNA gene V4 region from the preliminary targets in our database (S1 Doc). Curated databases were generated for each of the taxa in our preliminary target list using the performance metrics sensitivity, specificity, precision and negative predictive value as optimizing parameters. In other words, the bioinformatics pipeline was optimized to ensure that a positive result on the test truly means the target is present in the sample and a negative result is only obtained when no target is present in the sample. After optimizing the confusion matrices for all preliminary targets, 28 out of 46 targets passed our stringent threshold of 90% for each of the parameters (Fig 2). The resulting target list is composed of 5 known pathogens, 3 beneficial bacteria, and 20 additional microorganisms related to various gut afflictions, as well as commensal bacteria and one archeon. On average the sensitivity, specificity, precision and negative prediction value of the bacteria on our target list are 99.0%, 100%, 98.9% and 100%, for the species, and 97.4%, 100%, 98.5% and 100% for the genera, respectively.

**Fig 2.**
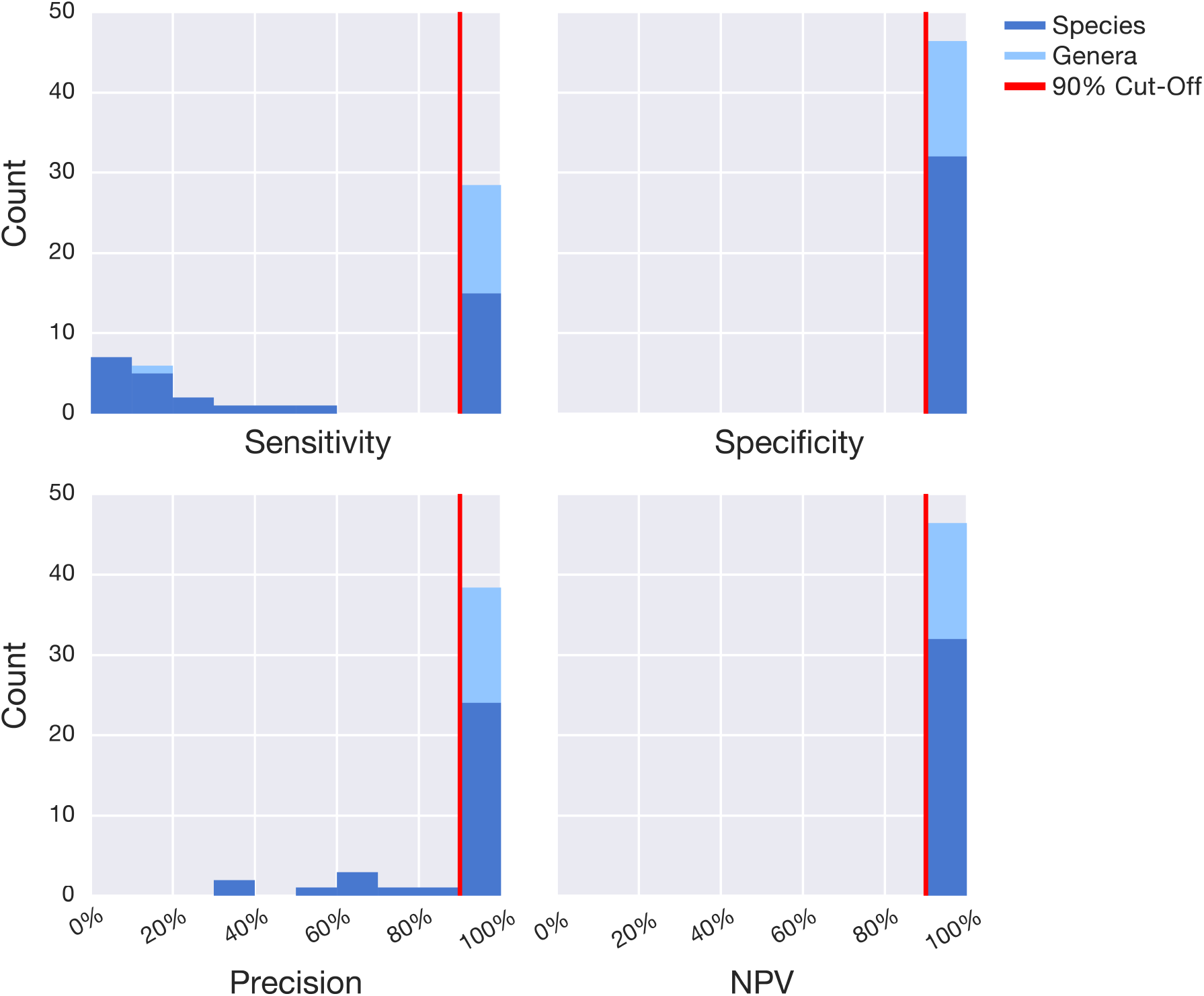
Bioinformatics target identification performance metrics.

The 46 preliminary targets identified from literature and available clinical tests are comprised of 15 genera and 31 species. To optimize the bioinformatics pipeline for accurate detection of the maximum number of targets, the following performance metrics were evaluated based on the number of true positives (TP), true negatives (TN), false positives (FP) and false negatives (FN) detected: specificity = TN / (TN + FP); sensitivity = TP / (TP + FN); precision = TP / (TP + FP); and negative predictive value (NPV) = TN / (TN + FN). After optimization, 28/46 preliminary targets passed our stringent threshold of 90% (red vertical line) for each of the parameters, resulting in the accurate detection of all genera (light blue) except for *Pseudoflavonifractor*, and 14/30 species.

### Reference ranges from a healthy cohort

Many clinically relevant microorganisms associated with health and disease are present at some level in the gut of healthy individuals. The clinical significance of microbiome test results is determined not only by the identity, but also the quantity of distinct species and genera within the context of a healthy reference range. To determine the healthy reference range for the 28 targets, we established a cohort of 897 samples from self-reported healthy individuals from the uBiome microbiome research study (manuscript in preparation). Microbiome data from this cohort were analyzed to determine the empirical reference ranges for the 14 species and 14 genera. For each of the 897 samples, we determined the relative abundance of each target within the microbial population. This analysis gave rise to a distribution of relative abundance for each target in the cohort (Fig 3). From these data, we defined a central 99% healthy range or confidence interval for each target. If the relative abundance of a target is outside of this healthy range within a sample, it would be considered a positive result.

Many of the targets show significant spread, emphasizing the importance of microbiome identification in the context of a reference range. For example, the pathogen *Peptoclostridium* 213 *difficile* is found in ~2% of the healthy cohort which shows that asymptomatic *P. difficile* colonization is not uncommon in healthy individuals [24]. Although all taxa are not found to be present in at least some of the healthy individuals, the maximum of the reference range can be quite high for some taxa (~63% for *Prevotella* and 49% for *Bifidobacterium*, for example). Two species are not represented at all within the central 99% of the healthy cohort: *Vibrio cholerae* and *Ruminococcus albus*. The absence of *V. cholerae* is suggestive of its pathogenic nature and its relatively rare occurrence in the developed world. However, *R. albus*, has previously been found to be enriched in healthy subjects in comparison to patients with Crohn's disease [25].

**Fig 3.**
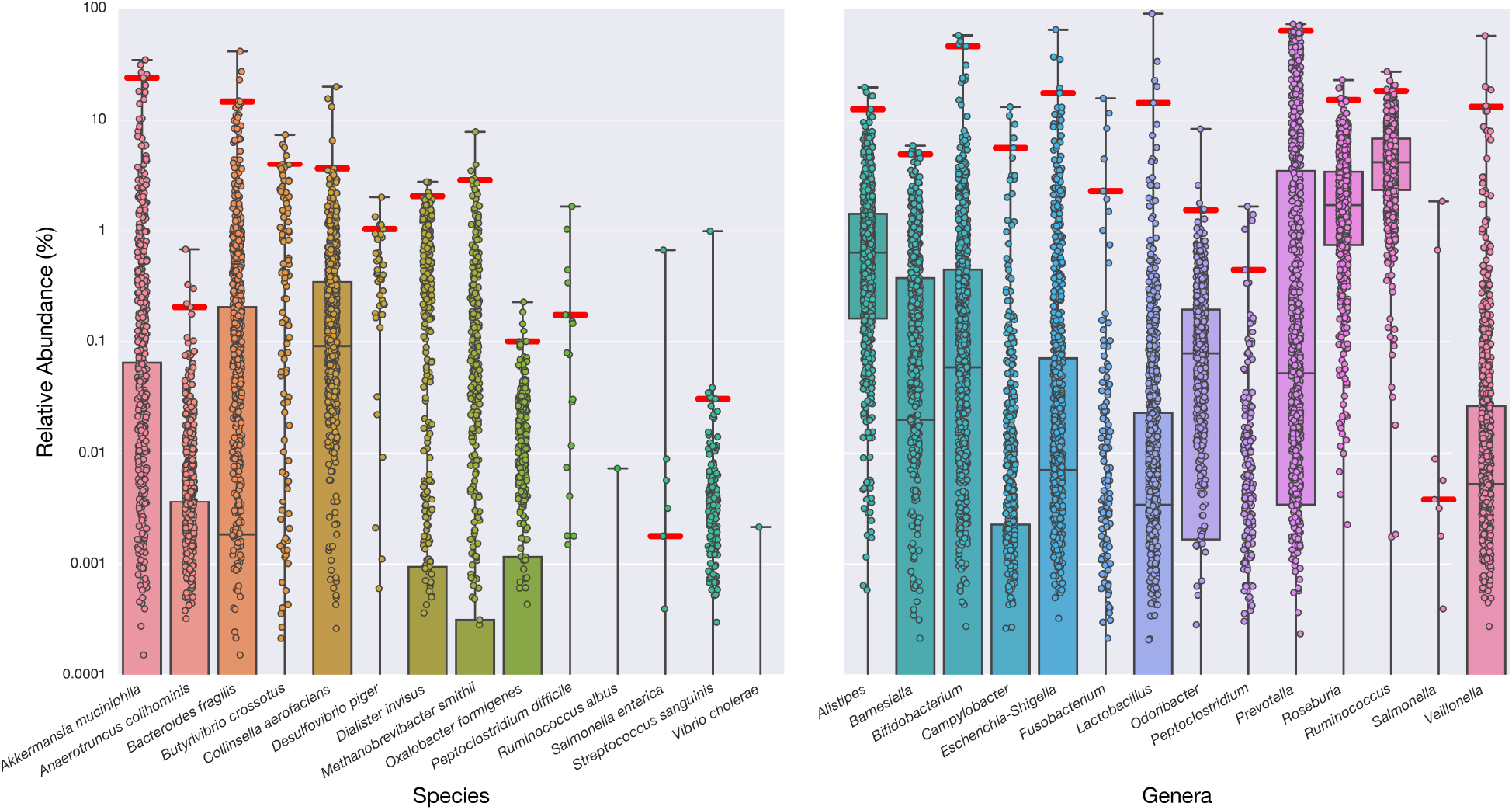
Reference ranges from a cohort of healthy individuals for 28 clinically relevant species and genera.

Healthy participant microbiome data were analyzed to determine the empirical reference ranges for each target. The boxplot displays the relative abundance for each of 897 self-reported healthy individuals, revealing the healthy ranges of abundance for the taxa in the test panel. The healthy distribution is used to define the 99% confidence interval (red whiskers). The box indicates the 25th–75th percentile, and the median coverage is indicated by a horizontal line in the box. Even in this healthy cohort, many of the bacteria that are associated with poor health conditions are present at some level. As most taxa are absent in a significant number of individuals most boxes expand to 0%, the healthy lower limit (not shown).

### Accurate detection of all 28 targets

To demonstrate our ability to accurately detect all the microorganisms in the clinical target list, we created representative synthetic double-stranded DNA (sDNA) gene blocks for each of the 28 targets (S2 Table). We analyzed the 28 targets as two sets of 14 distinct sDNA sequences. These sDNA sets were combined in specific proportions, resulting in five samples of increasing ratios for each target, 1:14,014, 1:1,414, 1:714 1:154 and ~1:14. We processed each sample using our clinical bioinformatics pipeline. Importantly, we accurately detected all targets at each ratio (Fig 4A), including the 1:14,014 ratio that is at our theoretical limit of detection, determined by our sequencing depth of 10,000 reads per sample (results shown for five species). Our ability to accurately determine the relative abundance of each target, even when it is present at exceedingly low levels, suggests that we can accurately detect each target within clinical samples and relate it to the healthy reference range to obtain a clinically informative result.

To further establish the clinical relevance of our pipeline, we tested 40 reference isolates from Luminex's xTAG Gastrointestinal Pathogen Panel. These verification samples comprise real or synthetic stool samples with live or recombinant material of known composition. Two of the 35 positive samples were excluded due to poor sequencing depth. The remaining 33 samples were positive for 1 of 8 different bacterial strains corresponding to 5 of our clinical targets: *Vibrio Cholerae* (5), *Salmonella enterica* (5), *Escherichia-Shigella* (13), *Campylobacter* (5) and *Peptoclostridium difficile* (5). All of the samples were correctly identified as having a relative abundance of the clinical target well above our defined healthy reference range (Fig 4B). Five samples containing *Yersina* were also tested as a negative control. Although *Yersinia* was included in our preliminary target list, it did not pass our stringent bioinformatics QC thresholds for accurate identification. As expected, the relative abundance of the 28 clinical targets was in the healthy range for the *Yersina* positive samples, as shown for *Escherichia–Shigella* (Fig 4B).

**Fig 4.**
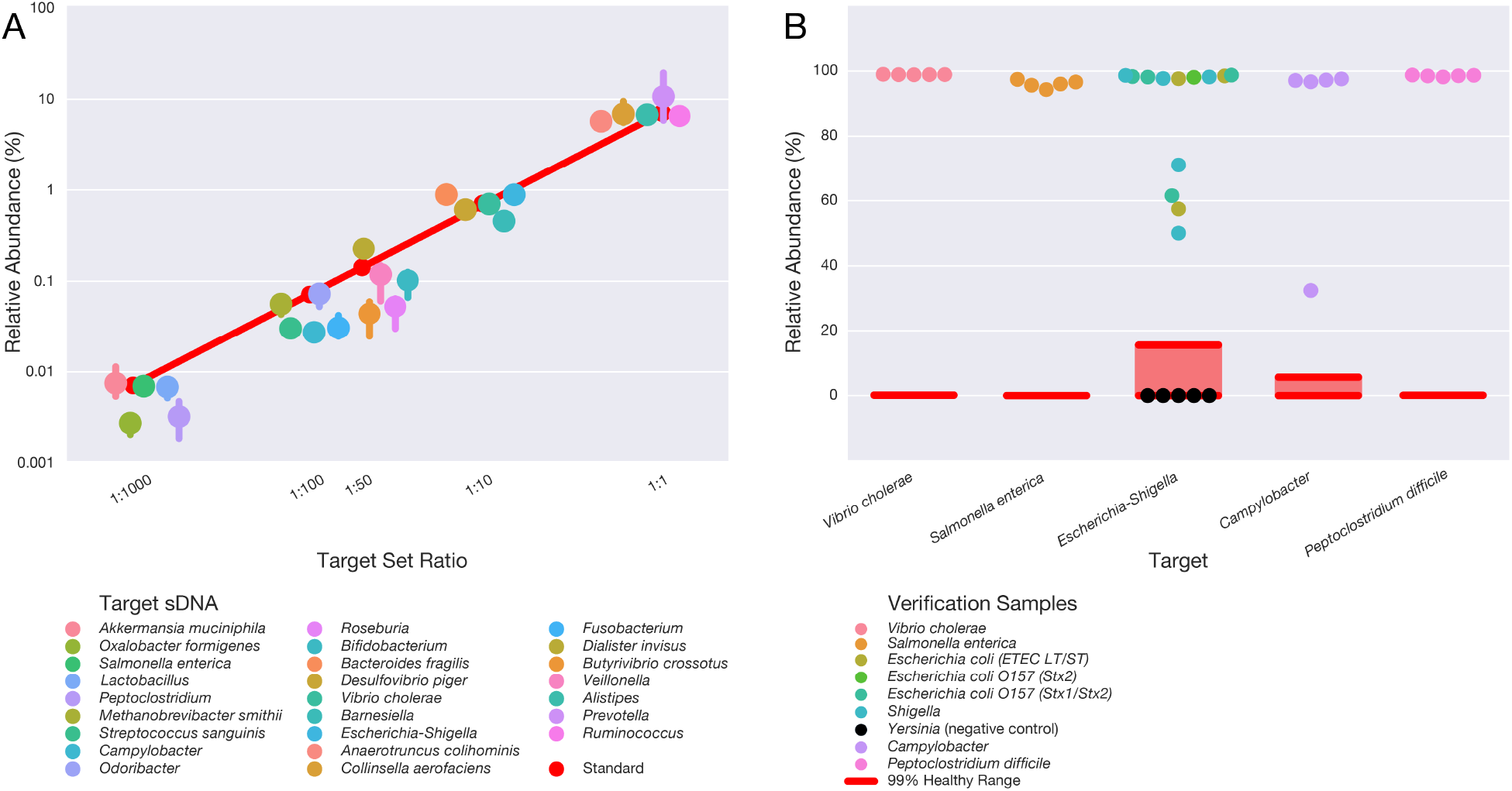
Experimental validation of the clinical 16S rRNA gene sequencing for the 28 targets on the test panel using synthetic DNA and verification samples.

A) Targets in the test panel are readily detected at various levels of relative abundance. Double-stranded DNA segments were designed to be representative for the V4 region of the 16S rRNA gene of each target species or genus. Two sets of 14 targets each were combined in 1:10, 1:50, 1:100 and 1:1000 ratios and vice versa, resulting in the following expected relative abundances of each target in the diluted set: 0.65%, 0.14%, 0.071% and 0.0071%. The abundance of the undiluted set in these ratios is plotted as 1:1 at an expected 7.1%. The expected abundances are plotted in red, while the average abundance of 5 representative targets is plotted with a confidence interval. The plot shows that targets are detected at different levels of relative abundance in the test panel within a high background of DNA.
B) Clinically relevant samples are accurately categorized using the bioinformatics pipeline. Verification samples containing real or synthetic stool samples positive for at least one control taxon from the target panel were tested using the bioinformatics pipeline. 33 distinct positive control samples, spanning 8 bacterial taxa were accurately identified at a level above the healthy range (red). All 33 control samples tested within the healthy range and thus were considered negative for the remainder of the taxa on our test panel. Five samples positive for *Yersinia*, a genus that is not present in our target list, were included as negative controls. These samples are visualized for the *Escherichia/Shigella* genus as they contained DNA for this taxon within the healthy range.

The accurate detection of a great number of microorganisms within a stool sample is critical to initiate the appropriate treatment in a clinical setting. Here we have shown that a test based on 16S rRNA gene sequencing can accurately detect and quantify clinically relevant levels of 28 target bacteria and archaea. We demonstrate that many prokaryotic targets identified from the literature as associated with human health can be consolidated in a single test, and further that relating the relative levels of bacteria and archaea to a healthy reference range enables the reporting of positive results only when clinically relevant.

The selection of microorganisms for this test panel was based on studies in medical journals and peer-reviewed articles. While all targets are relevant on their own, there is some overlap in the consolidated test. For example, while the *Salmonella* genus is unquestionably clinically relevant, testing for the genus when the test already includes the *Salmonella enterica* species might be redundant. The only other species of *Salmonella* is *Salmonella bongori*, a species that rarely infects humans and is mostly relevant to lizards [26].

While medical diagnosis has traditionally been focused on pathogens, research on the whole microbiome and its correlations with gut health continues to emerge. The test panel presented here reports on some microorganisms that are not usually interrogated in the clinic but provide additional insight into the overall gut health of a patient in a clinical setting. Because our detection method is based on DNA sequencing, the target panel can readily be expanded if new information about clinically important microorganisms arises. Because 16S rRNA gene sequencing identifies and quantifies the bacteria and archaea in a sample, relevant microbial metrics such as a microbiome diversity score can also be obtained, in addition to the information about individual targets, to provide a comprehensive overview of gastrointestinal health [27,28].

16S rRNA gene sequencing as a clinical diagnostic tool for gut-related conditions has many advantages over traditional culture-based techniques, including ease of sampling, scalability of the test, no need for human interpretation, and the ability to provide additional information about gut health. Thus, this method of detection for multiple clinically relevant microbial targets promises to have a real impact on patient diagnoses and treatment outcomes.

## Acknowledgements

This research did not receive any specific grant from funding agencies in the public, commercial, or not-for-profit sectors. We thank the uBiome lab team for sample processing, the bioinformatics team for data analysis, and all members of the uBiome team for helpful discussions. We thank Dr. Arthur Baca, Dr. Jonathan Eisen, Dr. Joe DeRisi, Dr. Alan Greene and Dr. Atul Butte for constructive input. We thank our scientific advisory board for their much-appreciated support. We thank Life Science Editors for editorial input. Finally, we especially want to thank all citizen scientist participants of the uBiome research study for their invaluable contributions.

## Supporting Information

### S1 Fig. Target sDNA Set Combinations

To demonstrate our ability to accurately detect all the taxa in the clinical target list, we created representative synthetic double-stranded DNA (sDNA) gene blocks for each of the 28 targets. The 28 targets were combined as two sets of 14 distinct sDNA sequences, target set A and target set B. Both of these sDNA sets were diluted and combined with the undiluted set in specific proportions (1:10, 1:50, 1:100 and 1:1000). Four dilutions of target set A were combined with the undiluted target set B and vice versa. The resulting ratio for each individual target in the diluted set is 1:154, 1:714, 1:1414 and 1:14014. The ratio of the individual targets in the undiluted set is 1:14.

### S1 Table. Bioinformatics Performance of the Preliminary Clinical Target List

The 46 targets identified from literature and available clinical tests comprise 15 genera and 31 species. The bioinformatics pipeline for accurate detection of the maximum number of targets is optimized based on the perfomrmance metrics Sensitivity, Specificity, Precision and Negative Predictive Value (NPV). The metrics are calculated based on the number of true positives (TP), true negatives (TN), false positives (FP) and false negatives (FN) as follows: specificity = TN / (TN + FP), sensitivity = TP / (TP + FN), precision = TP / (TP + FP) and negative predictive value (NPV) = TN / (TN + FN)

### S2 Table. Synthetic DNA Sequences (sDNA) for the Experimental Validation

The following representative synthetic double-stranded DNA (sDNA) gene blocks were synthesized for the 28 taxa in the target list. These sDNA sequences were run through the clinical pipeline to validate accurate and quantitative detection.

### S1 Doc. Extended Bioinformatics Methodology

